# CrowdVariant: a crowdsourcing approach to classify copy number variants

**DOI:** 10.1101/093526

**Authors:** Peyton Greenside, Justin M. Zook, Marc Salit, Ryan Poplin, Madeleine Cule, Mark DePristo

## Abstract

Copy number variants (CNVs) are an important type of genetic variation and play a causal role in many diseases. However, they are also notoriously difficult to identify accurately from next-generation sequencing (NGS) data. For larger CNVs, genotyping arrays provide reasonable benchmark data, but NGS allows us to assay a far larger number of small (< 10kbp) CNVs that are poorly captured by array-based methods. The lack of high quality benchmark callsets of small-scale CNVs has limited our ability to assess and improve CNV calling algorithms for NGS data. To address this issue we developed a crowdsourcing framework, called CrowdVariant, that leverages Google’s high-throughput crowdsourcing platform to create a high confidence set of copy number variants for NA24385 (NIST HG002/RM 8391), an Ashkenazim reference sample developed in partnership with the Genome In A Bottle Consortium. In a pilot study we show that crowdsourced classifications, even from non-experts, can be used to accurately assign copy number status to putative CNV calls and thereby identify a high-quality subset of these calls. We then scale our framework genome-wide to identify 1,781 high confidence CNVs, which multiple lines of evidence suggest are a substantial improvement over existing CNV callsets, and are likely to prove useful in benchmarking and improving CNV calling algorithms. Our crowdsourcing methodology may be a useful guide for other genomics applications.

## 1 Introduction

Copy number variation is a type of structural variation that involves large-scale duplications or deletions of parts of a chromosome. Copy number variants can have substantial effects on cell and organism phenotype and are associated with many kinds of human disease (Redon et al. 2006) (Feuk et al. 2006) (Sudmant et al. 2015). Numerous algorithms have been developed to characterize these variants from genotyping arrays and next-generation sequencing data (English et al. 2015) (Tattini et al. 2015) (Mills et al. 2011) (Kidd et al. 2008). However, these algorithms often have poor concordance on both the location and the type of copy number variant, particularly when characterizing small-scale (< 10kbp) CNVs (Scherer et al. 2007) (Pinto et al. 2011). One key challenge in further developing and assessing these algorithms is the lack of a large set of “gold standard” or reference copy number variants.

Crowdsourcing has been used successfully to obtain gold standard labels in projects such as Galaxy Zoo (Raddick et al. 2010), ClickWorkers (Ishikawa, ST and Gulick 2012), FoldIt (Cooper et al. 2010), and Zooniverse (Prather et al. 2013), but little investigation has been done to understand how crowdsourcing can be best utilized to analyze genomic variation. Basic questions include whether or not any domain expertise is truly needed, how large the crowd should be, and how to best display genetic variation to workers. We decided to investigate the use of crowdsourcing platforms to classify copy number variants and to address these basic questions. Google has developed the Crowd Compute platform to facilitate large-scale crowdsourcing problems, and we developed our framework with this platform to enable high throughput classifications.

## 2 Results

### 2.1 The CrowdVariant Framework

The CrowdVariant framework uses a crowdsourcing platform to display putative copy number variant sites to workers and aggregates classifications from a pool of workers to determine the copy number state. Using this framework, we first ran an experiment to compare non-expert and expert classifications on a pilot set of putative CNV sites and then expanded our classifications to curate a genome-wide set of high confidence CNVs [Figure 1].

**Figure 1.**
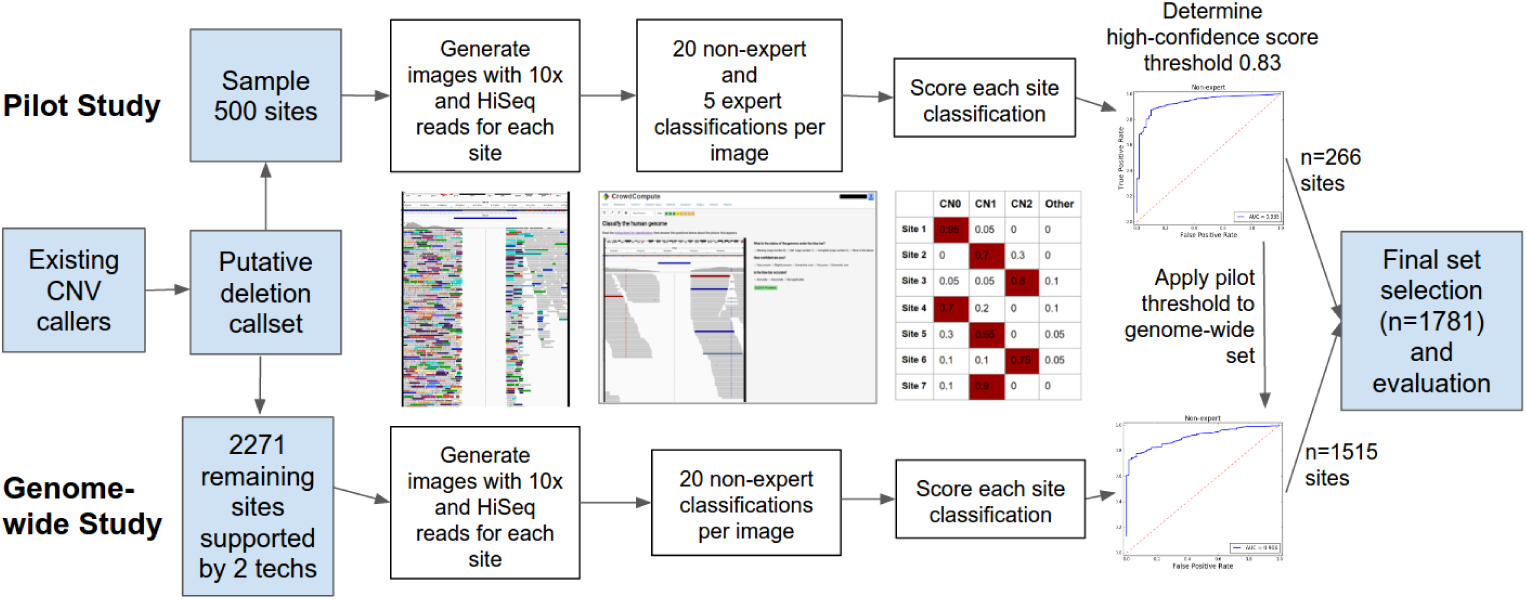
The experimental design was constructed to first evaluate a pilot set of sites with both experts and non-experts before applying the same framework to a genome-wide set of sites using non-experts only.

CrowdVariant displays pileup images of putative copy number variant sites using the Integrative Genomics Viewer (IGV), showing all reads aligned to the site and the flanking regions [Supplementary Figure 1] (Thorvaldsdóttir et al. 2013). Workers classify the site, assess break point accuracy and report their confidence based on seeing one image at a time.

We selected a set of 500 sites for the pilot phase of our study. We first called putative CNV sites using an ensemble approach from multiple technologies and algorithms [Supplementary Methods] (Abyzov et al. 2011) (Garrison and Marth 2012) (Mohiyuddin et al. 2015) (Hormozdiari et al. 2011) (Iqbal et al. 2012) (Mak et al. 2016) (Chaisson et al. 2014) (Nattestad and Schatz 2016) (Drmanac et al. 2010). We then randomly selected 500 sites ranging from 100bp to 3000bp with varying levels of support from existing algorithms [Supplementary Table 1].

We used aligned 10X Genomics (10X) and Illumina paired-end (Illumina) reads from the reference Ashkenazim trio made available by the Genome In A Bottle (GIAB)

Consortium (Zook et al. 2016). For each putative copy number variant site, we generated an image for each member of the trio using Illumina reads and an image for the son and the son’s reads separated by haplotype using 10X reads. Although workers potentially saw multiple images of the same site, we did not disclose to workers the experimental design, the technology, the individual or the site being shown.

In our pilot study, 20 non-experts each classified all 6 images for the 500 pilot sites. We launched a global recruitment for experts curators with over 110 individuals from several dozen institutions signing up to classify variants. The participation rate was highly variable with an average of 76 questions per expert [Supplementary Figure 2]. We ensured that all 6 images for at least 100 sites were classified by 5 experts each for comparison between experts and non-experts.

### 2.2 Non-experts can curate high quality copy number variants

Both experts and non-experts agreed on a consensus classification for the majority of sites [Supplementary Figure 3]. We visualized the responses for non-experts [Figure 2] and experts [Figure 3] by weighting each copy number classification and clustering workers and sites to reveal performance differences across sequencing platforms and individuals. We kept the identity of each non-expert worker separate, but we merged the expert answers into artificial workers 1 through 5 as experts did not answer enough questions individually to be meaningfully compared. For 86% of images, at least 70% of non-expert workers agreed on the classification, showing that non-experts can be trained to interpret copy number variants in a consistent manner [Supplementary Table 2]. Non-experts primarily had difficulty classifying haplotype images and systematically confused CN2s as CN1s for haplotype images only. Beyond these systematic errors, there were several non-experts that deviated from the majority either from lack of effort or understanding. Improving the documentation by showing more than 2 examples of each copy number type may further improve non-expert performance.

**Figure 2.**
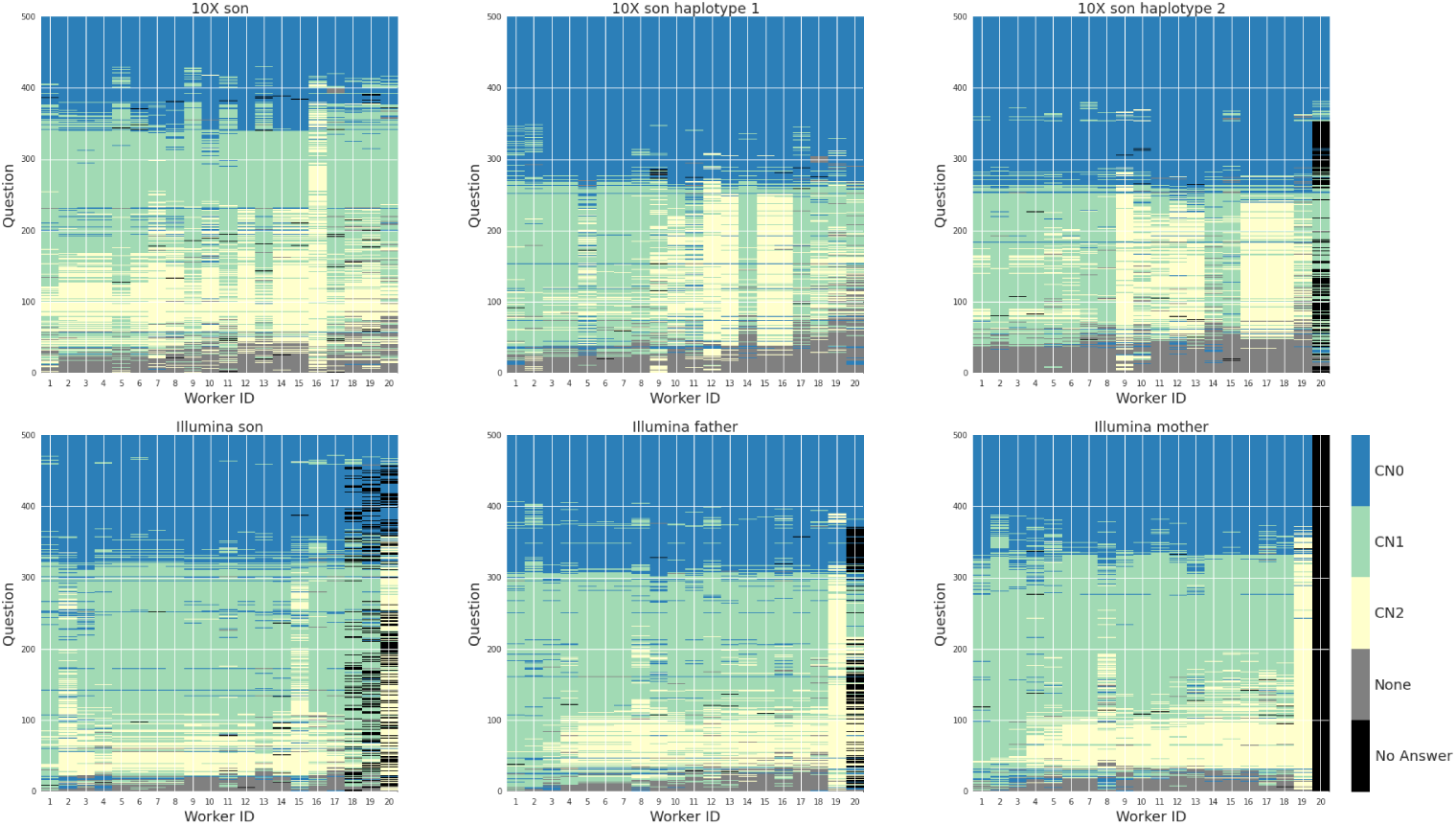
Non-expert classifications for 500 sites were color coded, weighted and clustered. Rows represent a question (i.e. an image of a putative site using a particular sequencing technology) and columns represent workers. Clockwise from top left: 10X son, 10X son haplotype 1 only, 10X son haplotype 2 only, Illumina mother, Illumina father, Illumina son.

**Figure 3.**
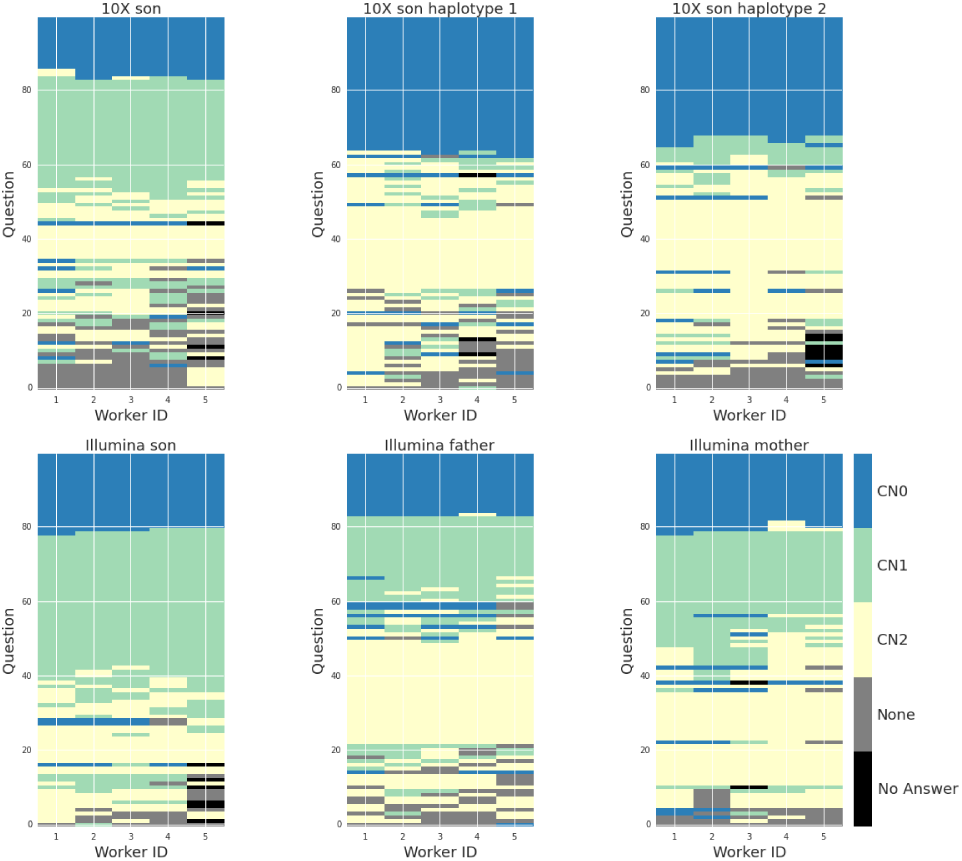
Expert classifications for 100 sites were color coded, weighted and clustered. Rows represent a question (i.e. an image of a site using a particular sequencing technology) and columns represent workers. Clockwise from top left: 10X son, 10X son haplotype 1 only, 10X son haplotype 2 only, Illumina mother, Illumina father, Illumina son.

Agreement among workers was used to assign a final classification and confidence score to each putative site. We defined the CrowdVariant score as the proportion of workers that voted in favor of the most popular classification (CN0/CN1/CN2/None of the Above), with higher scores reflecting more confident classifications. We incorporated worker classifications for all images of the same site, but classified each site for each individual in the trio independently. We counted all diploid classifications but only those haploid classifications where the pair of haplotype images was consistent with a diploid classification [Supplementary Methods]. We assign the most likely copy number state to each site by selecting the classification with the largest proportion of votes.

Non-experts performed similarly to experts when comparing the rate of Mendelian violations among the trio for each site [Supplementary Methods] [Table 1]. We found that 89% and 90% of all sites were classified without a Mendelian violation for experts and non-experts, respectively. The sites with Mendelian violations had lower scores and could largely be filtered out of the high quality set. The CrowdVariant scores discriminated Mendelian violations from plausible trio classifications with an AUC of 0.89 for non-experts and an AUC of 0.87 for experts [Supplementary Methods] [Supplementary Figure 4]. For comparison, we randomized all answers by re-sampling the entire worker by classification matrices for experts and non-experts and re-computed the rate of Mendelian violations [Supplementary Table 3]. The AUCs for expert and non-expert randomized answers were 0.47 and 0.50, respectively, and both 95% confidence intervals overlapped a random AUC of 0.5.

**Table 1.** Using metrics that rely on familial structure, non-experts perform at a similar level to experts.

| Metric \ Data Set | Expert | Non-expert |
| --- | --- | --- |
| Percent of sites without violation | 89/100 (89%) | 448/500 (90%) |
| ROC AUC | 0.87 | 0.89 |
| ROC AUC 95% confidence interval | [0.79, 0.95] | [0.86, 0.92] |
| Average violation probability | 0.15 | 0.14 |

**Table 2.** Reference and alternate read count cut-offs for each data set type. *For 10X, each haplotype was required to have at least 6 reference reads and <2 alternate reads, or at least 6 alternate reads and <2 reference reads. For heterozygous calls, one haplotype supported reference and the other alternate.

| Dataset | HomRef<br>minRef | HomRef<br>maxAlt | Het<br>minRef | Het<br>minAlt | HomVar<br>maxRef | HomVar<br>minAlt |
| --- | --- | --- | --- | --- | --- | --- |
| Illumina 150bp | 100 | 4 | 75 | 75 | 4 | 100 |
| Illumina 250bp | 20 | 2 | 10 | 10 | 2 | 20 |
| Illumina Mate-pair | 10 | 40 | 20 | 20 | 10 | 40 |
| 10X | 6* | 1* | * | * | 1* | 6* |
| PacBio | 10 | 0 | 5 | 5 | 0 | 10 |

We curated a high confidence set of CNVs for the son (NA24385) with high probability of correctness and no Mendelian violations [Supplementary Materials]. We initially intended to use self-reported confidence to filter lower quality classifications, but most non-experts consistently reported medium to high confidence despite minimal training [Supplementary Figure 5]. To avoid relying on self-reported confidence, we ranked all 500 sites by their CrowdVariant score and selected all sites with a higher score than the site with the first Mendelian violation. This violation occurred at score 0.83 and resulted in discarding approximately half of the sites for a total of 266 high confidence sites. The high confidence set of sites contains 122 CN0, 138 CN1, 5 CN2 and 1 “None of the above” classification. 252 out of 266 are supported by at least two other technologies. Importantly, for all sites in the high quality set that were classified by both experts and non-experts, there was 100% agreement (n=56 sites) between experts and non-experts.

### 2.3 CrowdVariant can classify CNVs with variable support or unclear breakpoints

CrowdVariant agrees with consensus classifications from existing algorithms, while also classifying variants that are challenging for existing algorithms. CrowdVariant scores assigned to each site are correlated with the number of technologies underlying the original calls [Figure 4]. CrowdVariant classifications also show strong agreement with svviz (Spies et al. 2015), a semi-automated visualization tool that determines whether each read supports the reference allele, alternate allele, or is ambiguous. We used a preliminary heuristic method to classify copy number variants based on the read counts supporting the reference and alternate alleles as determined by svviz for each dataset, and required agreement across all datasets that had clear support for a genotype [Supplementary Methods]. When comparing all high confidence classifications, agreement with svviz was 82%. CrowdVariant was able to resolve 26 sites that were uncertain for svviz, explaining part of the discrepancy. When we removed sites that were classified as “None of the Above” in CrowdVariant or uncertain in svviz, agreement was 91% between the two methods. Agreement with svviz also increased with the number of supporting technologies [Figure 5].

**Figure 4.**
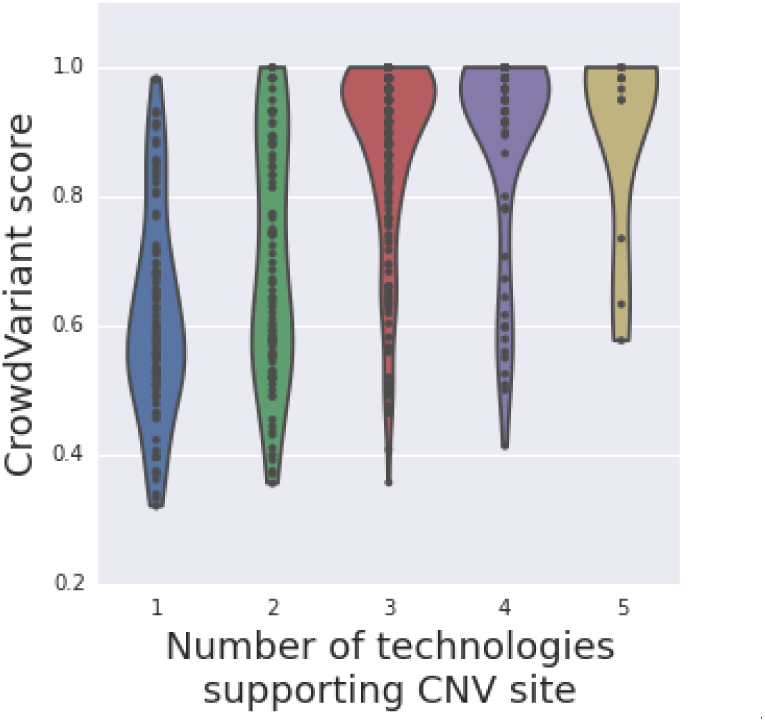
CrowdVariant scores determined by non-expert workers stratify by the number of supporting technologies from existing CNV callers.

**Figure 5.**
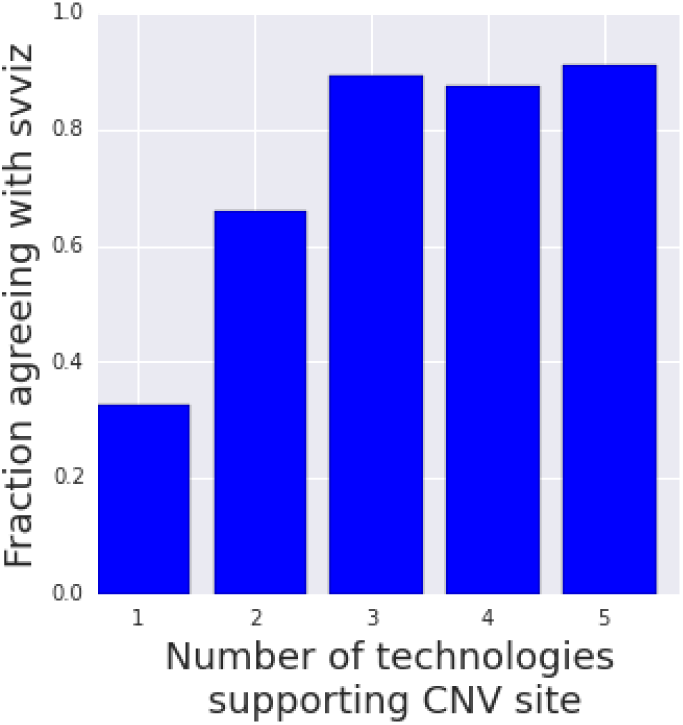
Agreement with svviz classifications for sites with varying support from orthogonal technologies. We only compare sites with CN0, CN1 or CN2 classifications from both methods.

The true power of incorporating many data types is clear when all 6 images of the same site are viewed together [Figure 6]. We find in multiple cases the crowd is able to resolve copy number state where other methods cannot, particularly when the boundary points are incorrect or ambiguous [Figure 7, Supplementary Figure 7]. While non-experts make some mistakes, we find that they do so in a consistent manner, such as mistaking a difficult-to-sequence region for a deletion, and they could likely be trained to recognize other features in the image that would clarify these mistakes. Phased data is particularly powerful for classifying heterozygous CNVs that are otherwise ambiguous and provides visual confirmation of the CrowdVariant results in conjunction with all other images for the site.

**Figure 6.**
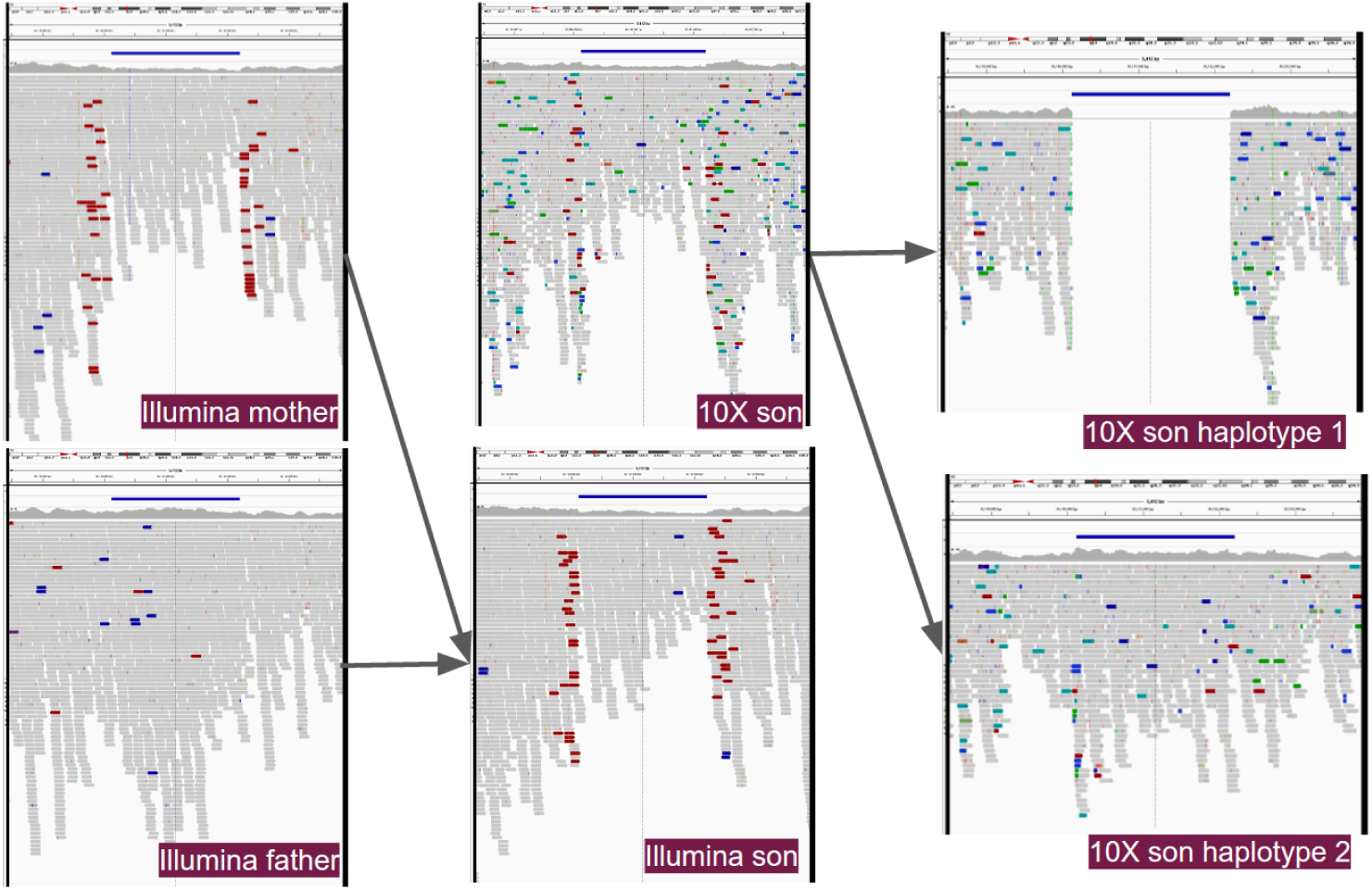
Viewing all image types together shows the power of combining familial and phasing information in different sequencing platforms. This variant (chr15:36160125-36162210) was classified as copy number 1 in the son with CrowdVariant score 1.0 and is part of the high quality set. The variant is visible in the mother, both diploid son images and one of the haplotype images. Clockwise from top left: Illumina mother, 10X son, 10X son haplotype 1, 10X son haplotype 2, Illumina son, Illumina father.

**Figure 7.**
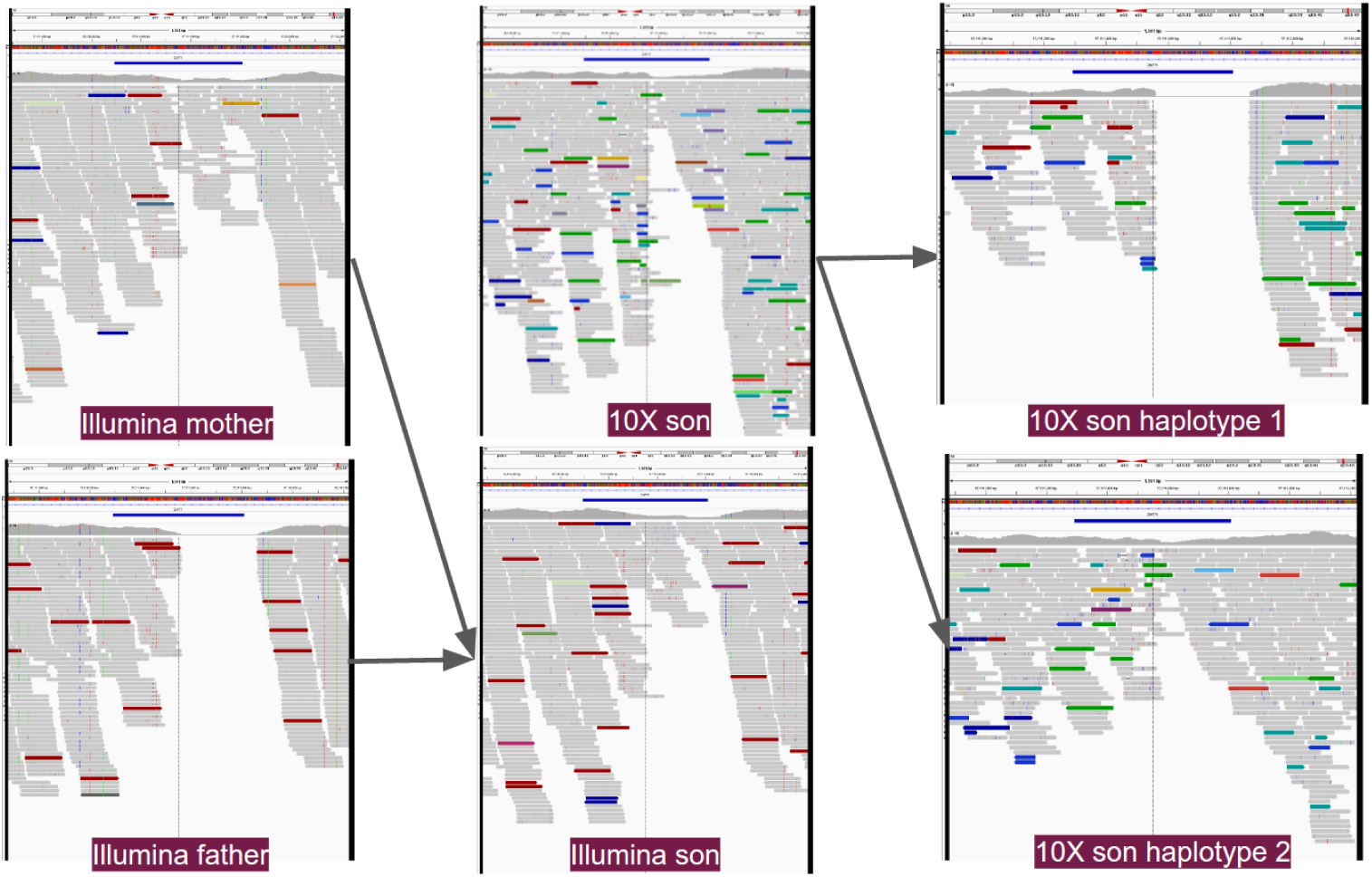
Viewing all image types together shows the power of combining familial and phasing information in different sequencing platforms. This variant (chr19:57111292-57111809) was classified as CN1 in the son with score 0.89 and is part of the high quality set. Svviz classified this example as CN2 due to the imprecise breakpoints. Clockwise from top left: Illumina mother, 10X son, 10X son haplotype 1, 10X son haplotype 2, Illumina son, Illumina father.

### 2.4 CrowdVariant can be used to curate a genome-wide high quality set of copy number variants

Having demonstrated that we can use non-expert workers to curate a high quality set of copy number variants, we expanded our classifications genome-wide. We took all putative CNV sites that were supported by GIAB callsets from at least 2 technologies and had not been classified in the pilot set (n=2271) and recruited 20 non-expert classifications for each site for all 6 image types. Due to the larger volume of images, not every worker classified all images in the genome-wide set. Consistent with the pilot study, we observed strong agreement among non-expert workers in the genome-wide set. Again, the primary inconsistencies were classifications for the haplotype images [Figure 8].

**Figure 8.**
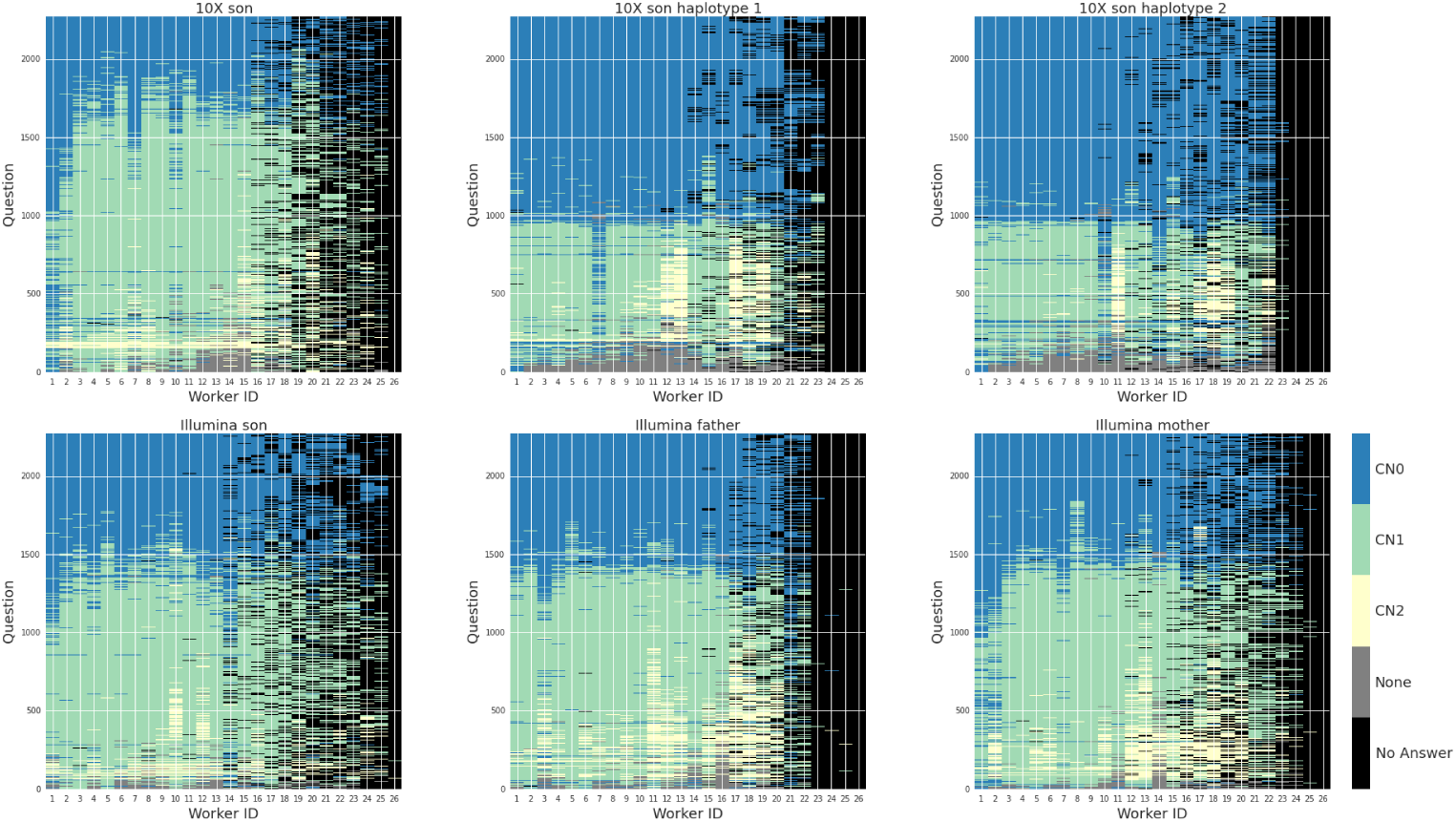
Non-expert classifications for genome-wide sites in Phase 3 were color coded, weighted and clustered. Rows represent a question (i.e. an image of a particular site using a particular sequencing technology) and columns represent workers. Clockwise from top left: 10X son, 10X son haplotype 1 only, 10X son haplotype 2 only, Illumina mother, Illumina father, Illumina son.

We scored each site by the proportion of workers voting for each classification and applied the threshold determined by the first 500 sites to curate high quality genome-wide classifications. This resulted in 1,515 new high confidence sites for the son (NA24385). The CrowdVariant scores for these sites correlate with the number of supporting technologies [Figure 9]. Likely due to requiring 2 supporting technologies, these sites were in even stronger agreement with svviz with 97.2% agreement among sites given CN0/CN1/CN2 classifications with both methods [Figure 10]. The high quality genome-wide set includes calls for 93 sites that svviz found uncertain. The additional genome-wide set includes 959 CN1, 552 CN0, 3 CN2 and 1 None of the Above. The CrowdVariant scores for the genome-wide set of CNVs also demonstrate similar concordance with orthogonal technologies [Figure 9] and classify Mendelian violations in the trio with auROC 0.94 [Supplementary Figure 8]. Above the threshold for high confidence determined from the pilot study, there was only one Mendelian violation in the genome-wide set occurring at a score of 0.94 [Supplementary Figure 9]. Combining with the 266 high quality sites from the pilot set, we finalized a set of 1,781 high confidence CNVs.

**Figure 9.**
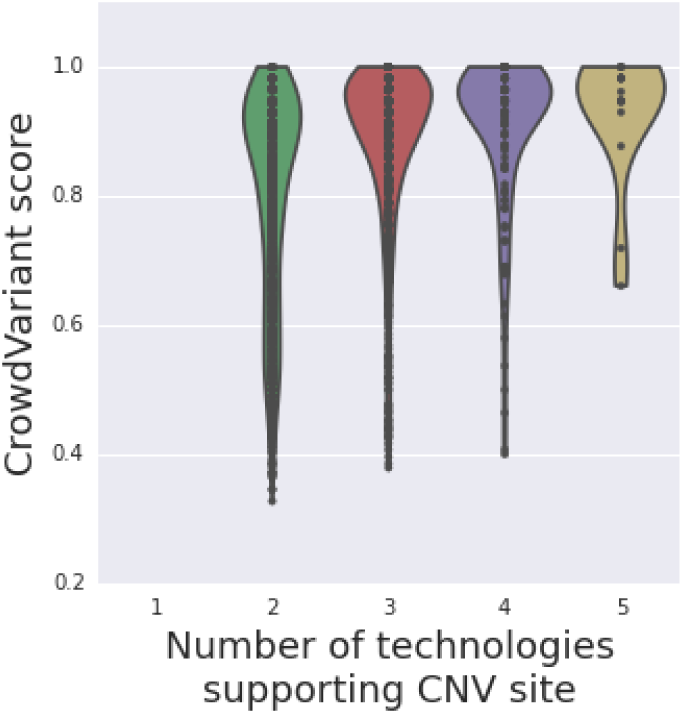
CrowdVariant scores for all genome-wide sites determined by nonexpert workers stratify by the number of supporting technologies from existing CNV callers.

**Figure 10.**
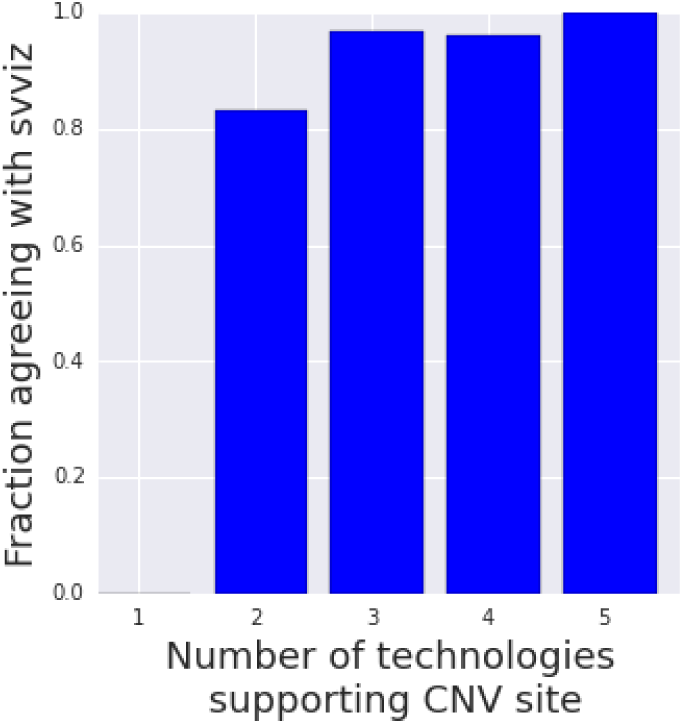
Agreement with svviz classifications for genome-wide sites with varying support from orthogonal technologies. We only compare sites with CN0, CN1 or CN2 classifications from both methods.

## 3 Discussion

We show that individuals with no background in genomics can be trained to accurately classify and thereby curate copy number variants. This is possible because, ultimately, the classification of CNVs based on images of aligned NGS reads is a pattern recognition problem, and even non-experts with limited training can excel at recognizing these patterns. As soliciting expert participation is prohibitively more difficult than non-expert participation (evident in the small amount of expert data we were able to collect), the ability to use non-experts enables crowdsourcing on a substantially larger scale. The larger scale afforded by non-expert workers allowed us to curate thousands of putative CNVs across the entire genome of a single individual from the Genome In A Bottle reference collection.

We are able to use non-expert classifications by using confidence scores to recognize the limit of their abilities. For many applications, such as deriving gold standard labels to improve machine learning methods, it is more critical to determine which classifications are trusted than to classify everything correctly. As machine learning approaches are increasingly adopted to solve genomic problems, crowdsourcing can provide an avenue to derive trusted training sets at high throughput for low cost.

While we have shown that crowdsourcing can be used to generate high confidence labels for CNVs, there are several limitations to our study. First, the set of CNVs we present is not a complete set for the GIAB Ashkenazim son (NA24385). Further, we only know that a CNV is segregating at the site, but we do not know its exact position or size. One broader limitation of crowdsourcing is that people can be consistent but wrong, however this limitation is shared by other approaches such as ensemble-based computational methods. In the current framework, our high confidence classifications are also enriched for sites that are overall easier to classify. However, there are many ways to increase confidence for more difficult questions by scaling the number of workers, augmenting training schemes, improving confidence metrics or considering alternative experimental designs such as those that incorporate both experts and non-experts depending on the particular question’s difficulty. Nevertheless, we are confident that our crowdsourced, genome-wide set of curated CNVs will prove valuable to methods developers working to improve their CNV calling algorithms.

Many possibilities exist for improving and expanding on this proof-of-concept study demonstrating the crowdsourcing curation of genomic variants. Incorporating images from additional technologies, such as long-read sequencing, could likely identify additional high confidence sites and remove some errors from using only short reads. Additional work might also use input from users about the precision of breakpoints. Other types of images could also be used, such as dot plots from assembly-assembly alignments and svviz images with reads mapped to reference and alternate alleles. These additional methods may help non-experts classify more difficult types of structural variants, like complex changes, insertions, inversions, and translocations, as well as variants in difficult, repetitive regions of the genome.

We use Google’s high throughput crowdsourcing platform, but as additional crowdsourcing platforms become available at low cost, soliciting participation from the crowd will become progressively easier. By using strategic experimental design, crowdsourcing can be a productive avenue to compete with and improve upon computational methods in difficult areas of genomics. Copy number variation, a domain where many experts still use manual inspection, is just one of these many areas. We provide a resource of high quality copy number variant classifications for a reference genome as a result of our study but ultimately see the potential expand far beyond these results.

## 4 Methods

### 4.1 Samples

We used the publicly available aligned reads for the Ashkenazim trio (NA24385/HG002, NA24149/HG003, NA24143/HG004) from the Genome In A Bottle (GIAB) Consortium. We selected the 300x Illumina HiSeq paired end sequencing data downsampled to 60x and the 10X Genomics Chromium dataset both for improved mapping using linked reads and the ability to separate the reads by haplotype. We used the pre-aligned bam files available from ftp://ftp-trace.ncbi.nlm.nih.gov/giab/ftp/data/AshkenazimTrio/.

### 4.2 Determination of Putative CNV sites

To form a preliminary integrated deletion callset for the Ashkenazim trio son (NA24385/HG002/RM 8391), we used callsets submitted to Genome In A Bottle (GIAB) from multiple technologies as of Aug 2016 for GRCh37. The calls are available at ftp://ftp-trace.ncbi.nlm.nih.gov/giab/ftp/data/AshkenazimTrio/analysis/NIST_DraftIntegratedDeletionsgt19bp_v0.1.7/. GIAB used a series of heuristics to integrate deletions from these callsets:

1. Find regions with deletion calls within 50bps of each other (i.e. sites are merged if they overlap by any amount or have <50bp separating them using bedtools merge −d 50).
2. Annotate each region with the fraction of bases covered by calls from each method.
3. Divide call regions into those that contain >25% tandem repeats (TR bed file with regions >200 bps in length containing TRs with >95% identity: https://github.com/ga4gh/benchmarking-tools/blob/master/resources/stratification-bed-files/LowComplexity/AllRepeats_gt200bp_gt95identity_merged.bed.gz). Remove all calls with TRs.
4. Find regions for which methods from at least 2 technologies have <20% difference in predicted size, treating all methods equally. The methods considered from each technology are as follows:
  - Illumina: spiral, cortex, commonlaw, MetaSV, Parliament/assembly, Parliament/assembly-force, CNVnator, GATK-HC, freebayes
  - PacBio: CSHL-assembly, sniffles, PBHoney-spots, PBHoney-tails, Parliament/pacbio, Parliament/pacbio-force, MultibreakSV, smrt-sv.dip, Assemblytics-Falcon, Assemblytics-MHAP
  - Complete Genomics: CG-SV, CG-CNV, CG-vcfBeta
  - Optical Mapping: BioNano, BioNanoHaplo (haplotype-aware)
5. Find the sensitivity of each method to these calls in the size bins:
  - 20-49bp
  - 50-100bp
  - 100-1000bp
  - 1000-3000bp
  - >3000bp
6. Filter all calls for which there is an overlapping call from any method that differs in size by >2x. Only include callsets if 75% of calls are <20% different from median size in the <3kb or >3kb size range (whichever the call falls in). This is intended to remove calls that may be more complex than just a simple deletion (e.g. a compound heterozygous event with deletions of different lengths or deletions of ambiguous length in repetitive regions).
7. Filter calls overlapping segmental duplications >10kb by 20% or more, or overlapping N’s in GRCh37 by any amount.
8. Use the breakpoints from the caller with the “best size accuracy” as described below.

Process for finding accuracy of predicted size from different techniques:

1. Find regions with support from >3 callers.
2. Find the difference between the size predicted by each caller and the median size of all callers with calls in each region.
3. Find the 75th percentile of this difference (i.e. 75% of calls have a predicted size < *x*% different from the median size).

Note that this is likely an overestimate of the size accuracy due to the limited number of callsets used and it does not account for any biases that might be present in all or most callsets.

### 4.3 Selection of CNV sites to show

For the pilot study we randomly sampled 500 CNV sites to use in our experiment to represent a diversity of putative site sizes and support. We binned all putative CNV sites into 5 size groups: 100-299bp, 300-399bp, 400-999bp, 1000-1999bp, 2000-2999bp. We sampled 100 sites from each size bin and within these size bins sampled 60 sites supported by 3 or more technologies, 20 supported by 2 technologies and 20 supported by 1 technology. We did no additional filtering of these randomly sampled sites. For the genome-wide set we selected all putative CNV sites supported by at least 2 technologies and within 100-2999bp that were not profiled in the pilot study.

### 4.4 Experimental Design

We represented each site with six different images in the CrowdVariant platform. We used 10X data for three images and Illumina paired-end data subsampled at 60x for the other three. The three images displaying 10X data showed the son (NA24385/HG002) as well as the same reads split by haplotype, each shown in a separate image. The three images using Illumina data displayed the family trio of son, father and mother (NA24385/HG002, NA24149/HG003, NA24143/HG004) at the same site.

We asked for 20 different non-expert workers to classify each image and 5 different experts to classify the same images. For the experts, we batched the 500 sites into 5 sets of 100 and prioritized classifying all 6 images for the first set of 100 sites before moving to the next 100 sites. Each worker could only classify the same site once, and the images were presented in a random order to each worker within each batch.

### 4.5 Image Generation

We used IGV to generate images for each site. We showed the site as well as 80% of the size of the region on either side for genomic context.

### 4.6 Participant Recruitment

Non-experts were recruited from Google’s CrowdCompute team. This group of workers is untrained in genomics applications. We recruited experts from the genomics community via email and social media. Workers answered as many questions as they were willing. Experts were determined by self-reporting some or substantial experience classifying CNVs.

### 4.7 Participant Instruction

Workers were shown an example pileup image highlighting three features useful to focus on: the putative CNV site location, the aligned reads and the overall distribution of reads over the reference genome. We showed two examples images each of copy number 0, copy number 1, and copy number 2. We also showed two example images each of accurate and inaccurate putative sites. The same training images were shown to both experts and non-experts, but the non-experts were shown modified language that did not contain technical terms (e.g. “bars” instead of “reads”). We did not bring additional attention to IGV features that may help image classification so that the documentation was the same for both groups, but experts may have made use of additional visual cues in the image given prior knowledge such as coloring for abnormally spaced reads or variant annotations.

### 4.8 Crowdsourcing Platform

We used Google’s Crowd Compute crowdsourcing platform. We developed a plugin that shows one image at a time along with a set of three questions. The image was not labeled with coordinates or any other identifying information beyond what is shown in IGV. Each worker answered 3 multiple choice questions about each image.

1. What is the status of the genome under the blue bar?
  Missing (copy number 0)
  Half (copy number 1)
  Complete (copy number 2)
  None of the above
2. How confident are you?
  Very unsure
  Slightly unsure
  Somewhat sure
  Very sure
  Extremely sure
3. Is the blue bar accurate?
  Accurate
  Inaccurate
  Not applicable

The answer for each question as well as the time to answer the questions was recorded. Workers were allowed to skip questions. We collected answers from the external community over 10 days.

### 4.9 Baseline voting model

We scored each possible copy number classification (CN0/CN1/CN2/None of the Above) at each site to determine the likely true copy number state as well as an associated level of confidence. We pursued several methods from a baseline voting scheme to methods that assess and re-weight some combination of each worker’s ability, the usefulness of each image type (i.e. sequencing platform) and the likelihood of mistaking one type of CNV for another. While the weighting framework appropriately minimizes the influence of ineffective workers and more difficult image types, we ultimately found that its benefits were minimal compared to a simple baseline voting scheme and we moved forward with a voting model for simplicity. In situations with larger numbers of more variable workers, the benefits of a more complex modeling scheme would likely be greater.

The baseline voting scheme scores each copy number state as the percentage of worker responses that voted in favor of that classification. For the diploid images, we included all worker classifications. However, the haplotype image classifications were only counted if the classifications for the paired haplotype images were genetically plausible. We initially allowed haploid CN0 and CN0 to count for diploid CN0, haploid CN0 and CN2 to count for diploid CN1 and haploid CN2 and CN2 to count for diploid CN2. Only haploid “None of the Above” and “None of the Above” counted toward “None of the Above”. All other configurations (e.g. haploid CN0 and “None of the Above”) were omitted from the vote. However, we observed a small boost in performance by allowing non-expert CN1 to count for CN2 in the haploid data since these errors were made systematically. Thus in the final model we also allowed haploid CN0 and CN1 to count for diploid CN1 and haploid CN1 and CN1 to count for diploid CN2. Using this voting model, we calculated for each site and for each individual in the trio a score for CN0, CN1, CN2 and “None of the Above” that sum to 1. The CrowdVariant score is the max of these scores.

### 4.10 Weighted Framework

We designed a model with three parameter sets based on our observations of the data. We note that despite using the same documentation, some non-experts appear to be more accurate than others. In addition, some of the images - whether due to sequencing depth, alignment quality or other factors - make it easier to determine the true copy number state from a pileup image. Finally, upon observing that workers can systematically mistake one type of copy number event for another, we note that we should be able to compensate for these types of systematic errors. Given these considerations, we define the score of each site to be a function of all the classifications for that site weighted by the worker ability and image usefulness and subject to systematic errors indicated by a confusion matrix mapping each classification to its likely intended classification.

Because we model latent diploid copy number state, we need to map the haplotype classifications to the diploid latent states that they support. We introduce the genetic function *G* in order to achieve that mapping. For diploid states, the genetic function returns the same state. For haploid data considered together, the genetic function will return the diploid state the data supports. Two haploid CN0s support diploid CN0 while haploid CN0 and CN2 support diploid CN1 and haploid CN2s support diploid CN2. When deciding how to translate haploid classifications with uncertainty (None of the Above answers) or discordant haplotypes (such as CN1 and CN1), we allowed two haploid “None of the Above” to contribute to “None of the Above” but otherwise prevent any discordant haplotype information from contributing to a classification. With our three parameter sets, genetic function and observed and latent data, we describe the model updates below:

- Iterables: sites *S*, individuals *I*, genotypes *G*, workers *W*, image types *M*
- X: observed classifications for each site by each worker
  indexed 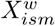
- C: latent true CNV state for each site
  indexed 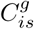
- *θ*: parameters
  W: worker ability
    indexed *W_w_*
  A: image usefulness
    indexed *A_m_*
  M: confusion matrices
    indexed 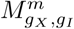

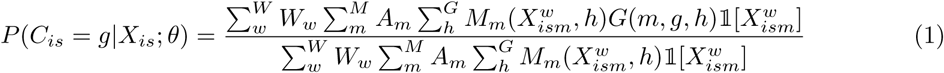

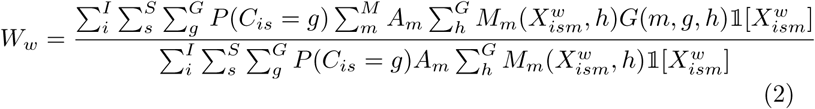

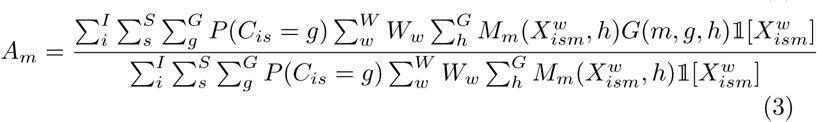

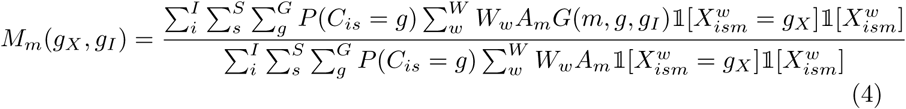

Noting that each parameter as well as our latent copy number state variables are defined in terms of one another, we use an iterative procedure to update each parameter and latent states one at a time. We initialize with equal worker abilities and equal image usefulness and confusion matrices as identity matrices.

We tried several iterations of the model updating all the parameters separately and in various combinations together. Upon updating all parameters, we discovered quickly that updating the confusion matrix based on highly variable classification data leads to severe over fitting of the data. We thus fixed the confusion matrices as the identity matrices for diploid classifications and to fix the confusion matrix for haploid sites allowing any CN1 to count toward a CN2. This means that haploid CN2/CN0 and haploid CN1/CN0 both count toward diploid CN1 and haploid CN2/CN2 and haploid CN1/CN2 both count toward diploid CN2.

### 4.11 Metrics for evaluation

There is no gold standard data available for all putative sites we evaluated. As a result, we rely on three metrics based on the familial structure of the 3 trio individuals to assess accuracy in addition to comparing to existing methods. First, we determined whether any Mendelian violations were made using the assigned most likely classifications for each site for each individual. For example, given a copy number 0 in the son, a copy number of 2 from either parent would be considered a Mendelian violation. For each site, we convert scores to hard classifications, determine whether a Mendelian violation was made and report the percent of sites that do not result in Mendelian violations. This metric has the caveat that some configurations, such as a CN1 in the son, are less likely to be caught if incorrect. However, the strong inverse correlation between our scores and Mendelian violations suggests it is sufficiently sensitive to evaluate our classifications as a whole.

In addition to the percentage of Mendelian violations, we assessed how well we could identify high confidence genetically plausible sites. We ranked the sites based on their CrowdVariant score (1 being most confident and 0.25 being least confident) and calculated the area under the ROC curve to assess how well our calculated scores discriminate genetically plausible from implausible sites. We also evaluated the uncertainty for the AUC using DeLong’s method to estimate 95% confidence intervals (Robin et al. 2011) (DeLong et al. 1988).

For the third metric, instead of relying on the most likely copy number status for each site, we calculated the total probability of a Mendelian violation. For every violating configuration we added the product of the three probabilities that the members of the trio took that genetic configuration. This total genetic error allows more nuance in determining whether our improvements in estimated probabilities are more plausible even if the most likely site does not change. The total probability of a Mendelian violation was 14% for all 500 non-expert classified sites and 15% for the 100 expert classified sites [Supplementary Figure 6].

We compute each evaluation metric separately for the expert and non-expert data. Visualization of the ROC curve indicates that the small samples size of questions for the experts may make the AUC less reliable than for the non-experts, motivating our incorporation of 95% confidence intervals. For comparison, we randomized all answers by re-sampling the entire worker by classification matrices for both experts and non-experts and re-computed the same metrics. The rate of genetic plausibility was around 54% for randomized expert answers and 73% for randomized non-expert answers. The higher number of plausible sites for non-experts is likely due to the increased number of CN1 and smaller number of CN2 classifications they gave, which have more genetically plausible configurations.

### 4.12 Comparison to svviz

In order to assess evidence for each of the calls in this work and assign preliminary genotypes, we used svviz (Spies et al. 2015) to determine whether reads from five data sets support the reference allele, the alternate allele, or were ambiguous for each member of the trio. The five data sets used were (Zook et al. 2016):

- ∼300× 2×150bp Illumina paired end sequencing
- ∼45× 2×250bp Illumina paired end sequencing
- ∼10-15× 2×100bp Illumina mate-pair sequencing with ∼6kb insert size
- ∼25-60× 10X Genomics Chromium sequencing, with reads separated by haplotype using bamtools filter
- ∼30-70× PacBio Sequencing

We used the “ref_count” and “alt_count” outputs from svviz batch mode, which correspond to the numbers of reads unambiguously supporting the reference and alternate alleles, respectively. For the results of each dataset from all genomes, we visually examined the density of sites in a plot of log(alt_count) vs log(reLcount), as well as in histograms of alt_count and reLcount. Based on the density of sites in these plots, we chose cut-offs for alt_count and reLcount for each dataset to define likely homozygous reference (CN2), heterozygous (CN1), and homozygous variant (CN0) sites. If the site had few reads assigned to reference or alternate, or it it did not fall within the bulk of sites with any genotype, then it was assigned an ambiguous genotype for that dataset.

A consensus genotype was then assigned if all datasets with an assigned genotype of CN0, CN1, or CN2 had the same genotype for that site. Sites with discordant genotype or with all ambiguous genotypes were given an uncertain consensus genotype. The uncertain classification in svviz is not directly comparable to “None of the Above” responses in CrowdVariant as some participants may have indicated a CN0/1/2 classification but marked low confidence to specify uncertainty while some may have specified “None of the Above.”

When we computed the rate of Mendelian violations among the svviz classifications we found 489 out of 500 (97.8%) without Mendelian violations. One reason that svviz classifications have higher concordance is because there are larger number of “None of the Above” classifications, which cannot be used to detect Mendelian violations. In addition, we observe the same decreased rates of Mendelian violations when calculating site probabilities using only Illumina data (493 out of 500 (98.6%)). Based on careful examination of the data, we believe this is because images of the same type often show the same kinds of biases consistently among a family trio. A consistent bias from one platform can lead to increased concordance, but not necessarily an increase in ground truth accuracy. Combining high quality data from diverse sources should improve our best estimate of the truth. Further we are not using genetic concordance to capture ground truth for individual sites, but instead to determine a threshold at which we have high confidence in crowdsourced classifications and to compare experts and non-experts. When we remove all sites for which svviz reported an uncertain genotype for any individual, we observe a Mendelian concordance rate of 398/398 (100%). When we similarly remove any “None of the Above” classifications from CrowdVariant responses, we observe a Mendelian concordance rate of 423/458 (92.4%), but note that an overall larger number of sites are classified.

For more information, see ftp://ftp-trace.ncbi.nlm.nih.gov/giab/ftp/data/AshkenazimTrio/analysis/NIST_DraftIntegratedDeletionsgt19bp_v0.1.7/SVSummary_v0.1.7_TRs.html.

### 4.13 High confidence sites

To determine high confidence sites we again ranked all of the 500 sites from the pilot study by the maximal score across all four classifications (CN0/CN1/CN2/None of the Above) for the son and identified the score at which the first Mendelian violation occurred. We determined all sites above this threshold as high confidence.

### 4.14 Genome-wide analysis

We selected all putative CNV sites supported by at least 2 technologies and not profiled in the first 500 sites (n=2271). We again recruited 20 classifications for all remaining sites by non-experts and used a baseline voting model to calculate final scores for each site. We used the threshold as determined in the pilot study to determine genome-wide high confidence sites. Using the same methods as described above for the pilot study, we evaluated the Mendelian violation rate, agreement with supporting technologies, and agreement with svviz.

## Data Access

All data is available in Supplementary Materials. We provide the scores for each putative copy number variant site and label the high quality sites. All raw worker answers for both non-experts and experts are available as well.

## Acknowledgments

We would like to thank Igor Karpov for his help working with the Crowd Compute infrastructure. We would like to thank Laura Paragano for her help in developing training materials. We are grateful to all participants who classified variants. We would like to thank Jonathan Bingham for his help in making this study possible. We would also like to thank the Verily analysis team for their feedback on the project.

## Author Contributions

P.G., J.Z., M.C., R.P., and M.D. designed the study. P.G. and M.D. developed the copy number variant crowdsourcing platform. J.Z. generated the results for orthogonal tools and determined putative CNV sites. P.G. developed the training and recruitment materials. P.G. performed analysis and modeling of the results. P.G., J.Z., M.C., M.S., and M.D. wrote the manuscript.

## Disclosure Declaration

M.D., R.P., M.C., and P.G. are employees of Verily Life Sciences, a for-profit corporation and affiliate of Google Inc. R.P. and M.D. are employees of Google Inc. Verily developed the CrowdVariant platform discussed in this paper, and could derive direct or indirect commercial benefit from positive research results pertaining to CrowdVariant.

## 5 Supplementary Figures

**Supplementary Figure 1.**
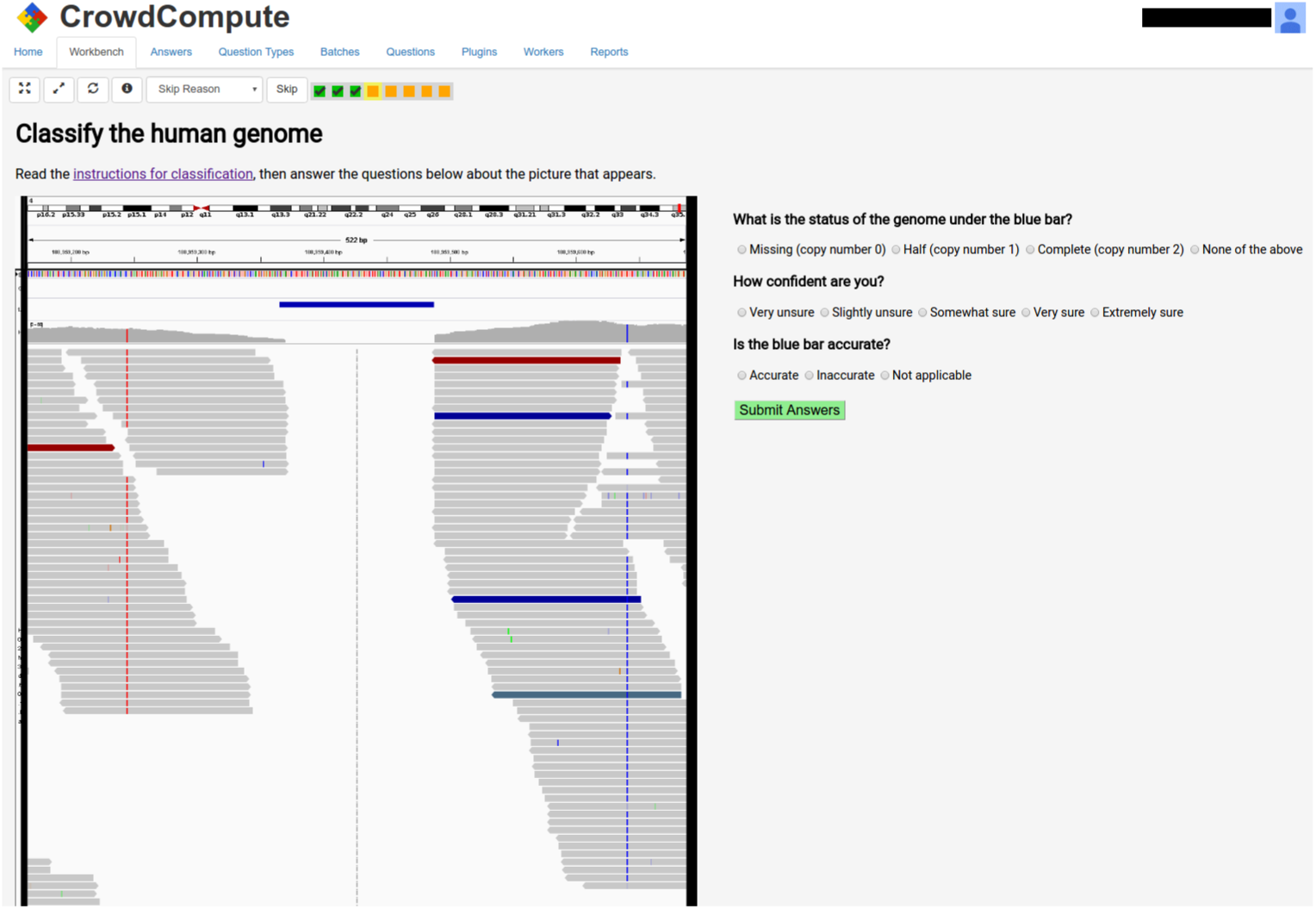
A screenshot of the CrowdVariant crowdsourcing platform.

**Supplementary Table 1.**
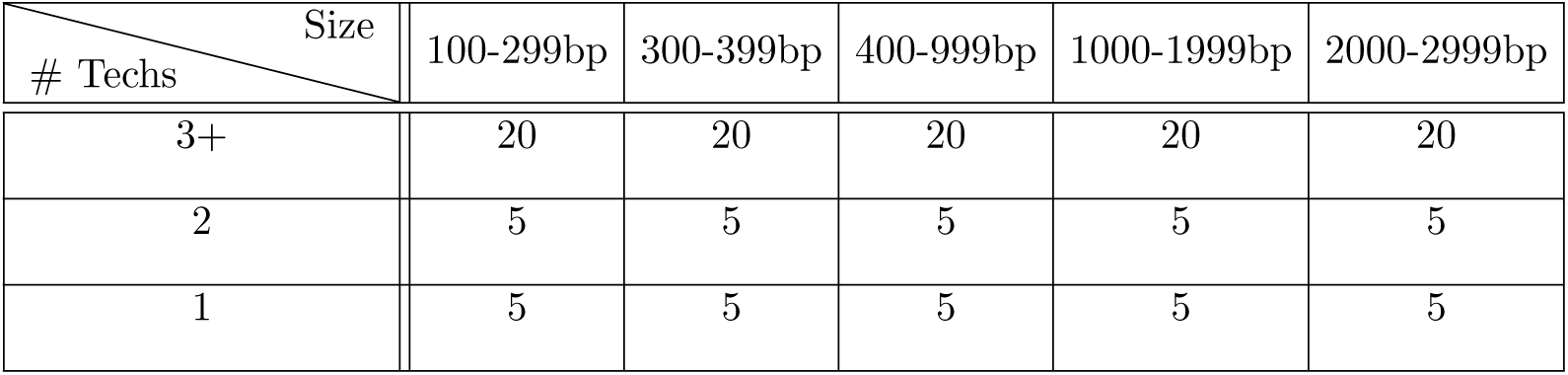
Number of sites sampled for each region size window and number of sequencing technologies supporting the putative CNV site.

**Supplementary Figure 2.**
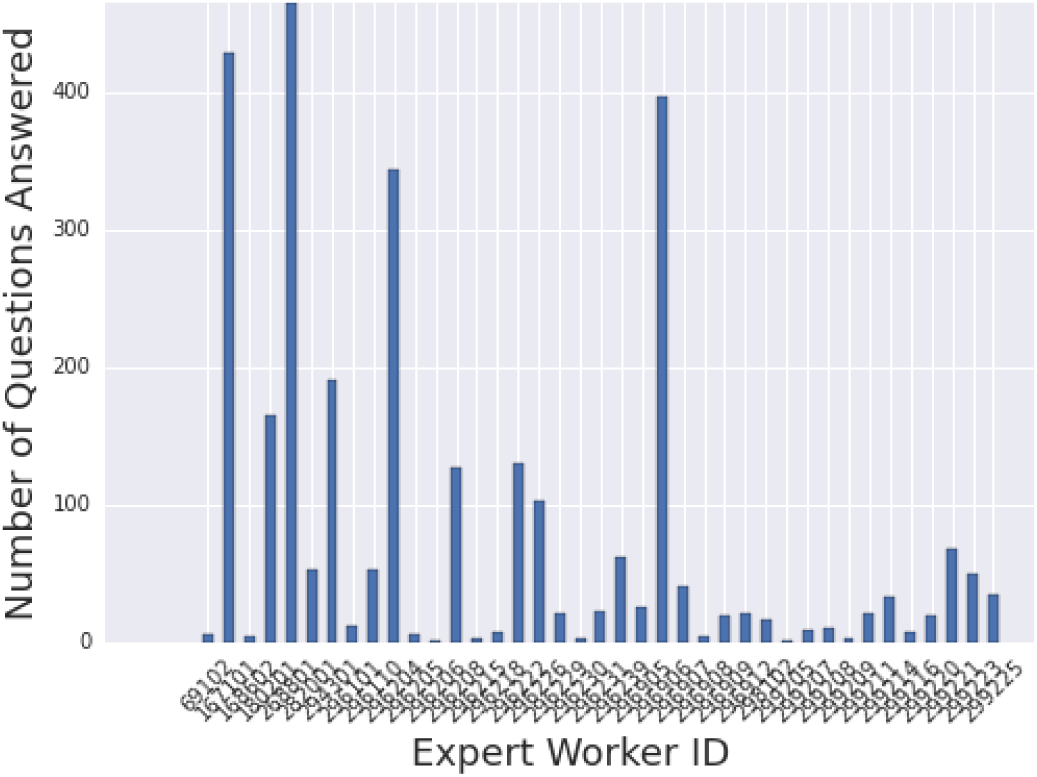
The number of questions answered by each expert worker ranged from several to several hundred.

**Supplementary Table 2.**
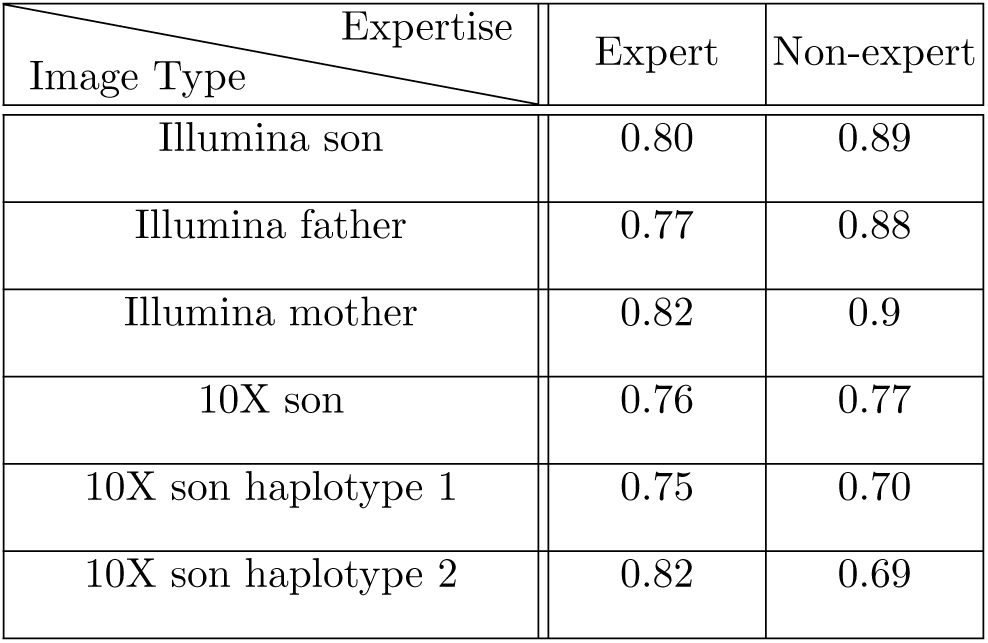
Mean number of sites with a concordant answer among workers for each image type. We set the concordance level at 70% of responses in agreement and computed concordance rates above this threshold for every image type separately for both expert and non-expert classifications.

| Image Type \ Expertise | Expertise |  |
| --- | --- | --- |
|  | Expert | Non-expert |
| Illumina son | 0.80 | 0.89 |
| Illumina father | 0.77 | 0.88 |
| Illumina mother | 0.82 | 0.9 |
| 10X son | 0.76 | 0.77 |
| 10X son haplotype 1 | 0.75 | 0.70 |
| 10X son haplotype 2 | 0.82 | 0.69 |

**Supplementary Table 3.**
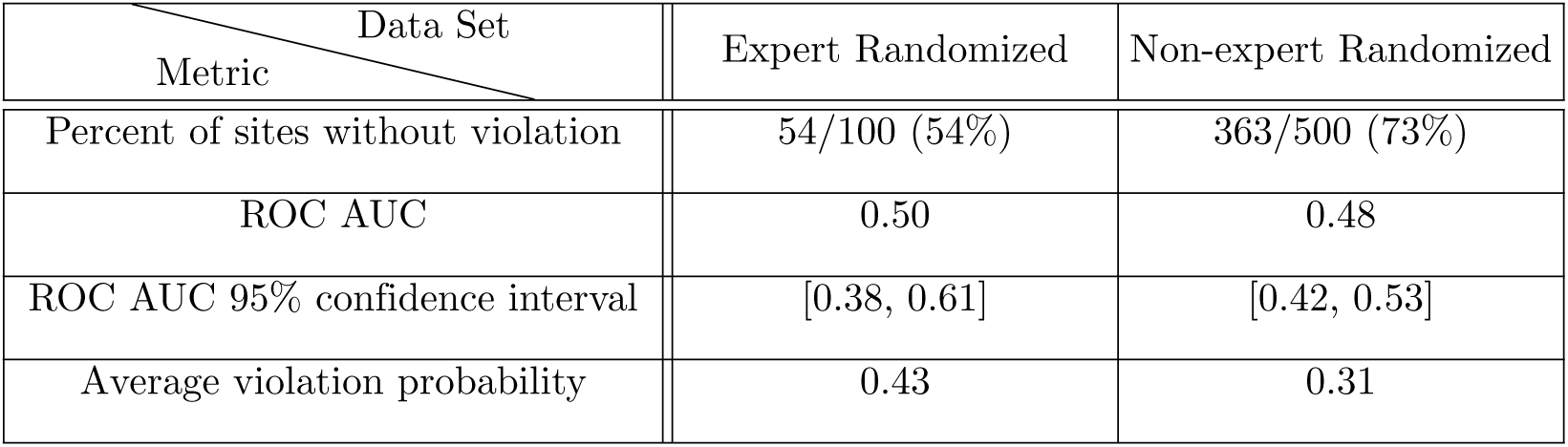
Randomized answers a give lower bound on violation metrics. We randomized the answer by worker matrix and re-computed evaluation metrics. Both AUC confidence intervals overlap a random AUC of 0.5.

**Supplementary Figure 3.**
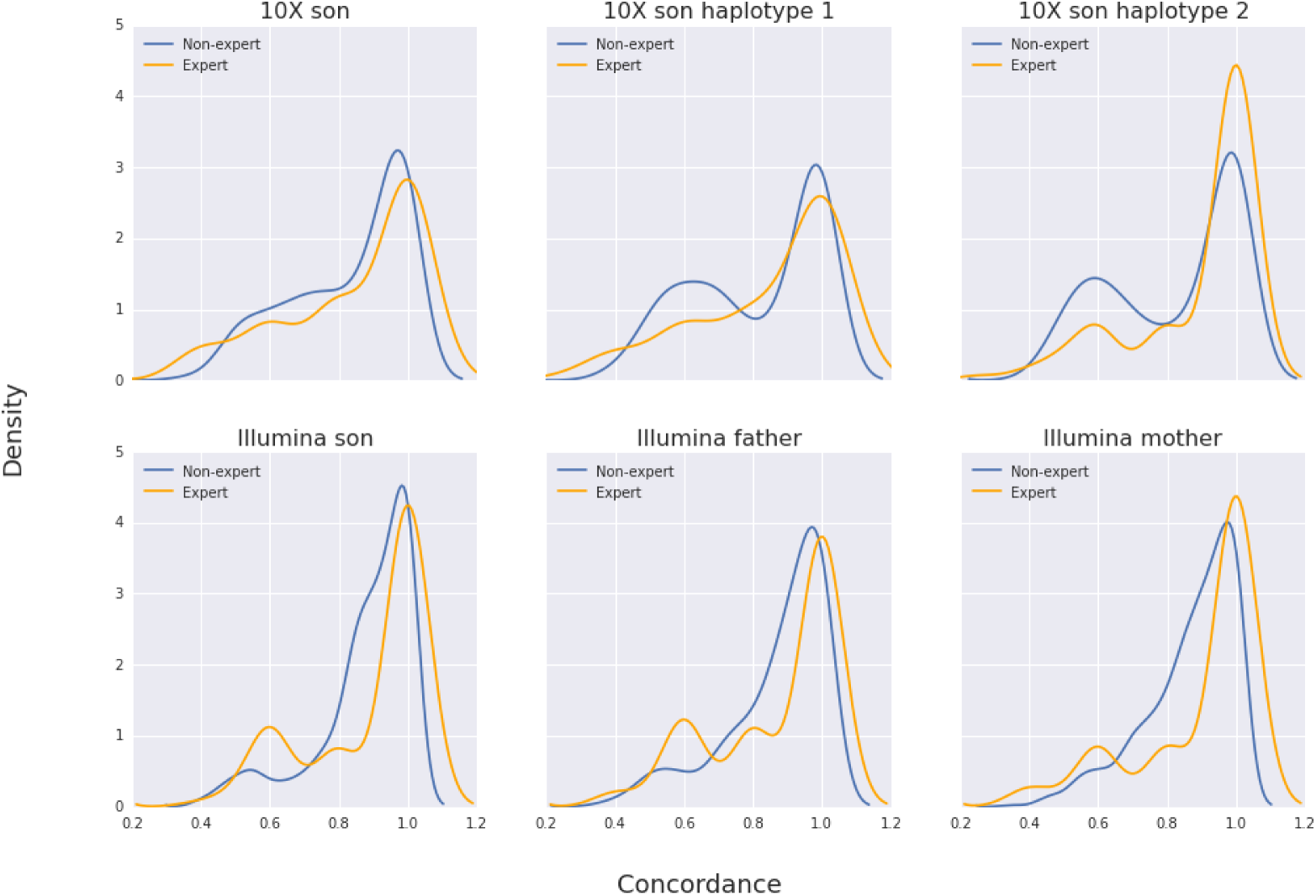
Kernel density of percent agreement on most common classification for each image type. Experts and non-experts show similar levels agreement.

**Supplementary Figure 4.**
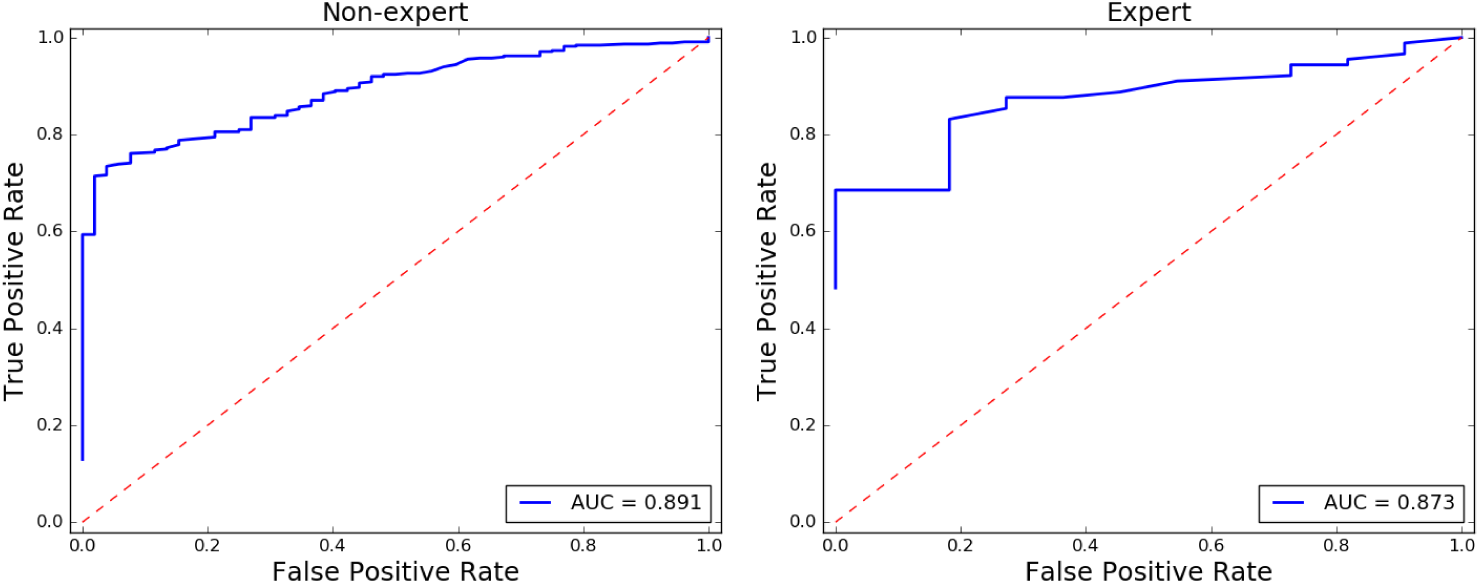
Non-expert (left) and expert (right) ROC curves discriminating Mendelian violations (1=no violation, 0=violation) sites by the CrowdVariant score.

**Supplementary Figure 5.**
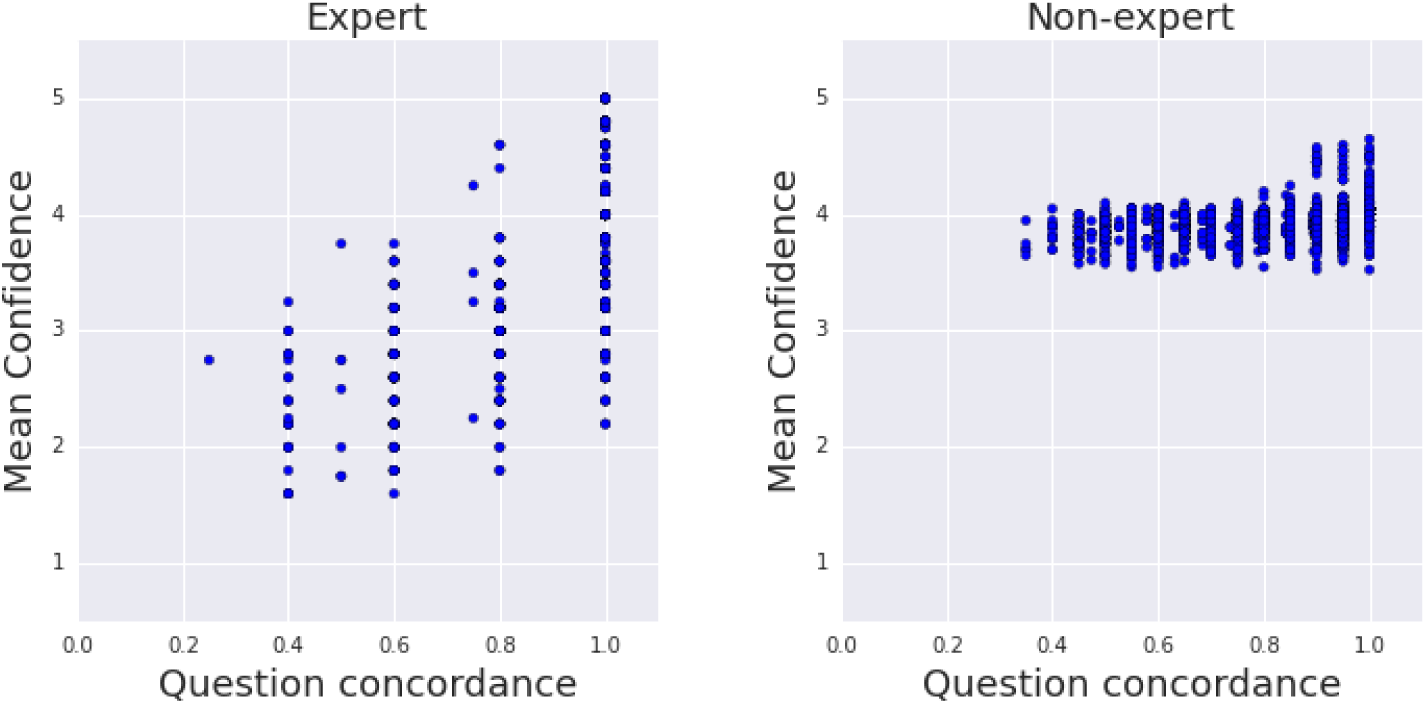
Mean reported confidence scores for each question as a function of concordance (percent agreeing with most common answer) for experts [left] and non-experts [right]. Non-expert reported confidence does not display enough variation to sufficiently filter answers based on confidence alone.

**Supplementary Figure 6.**
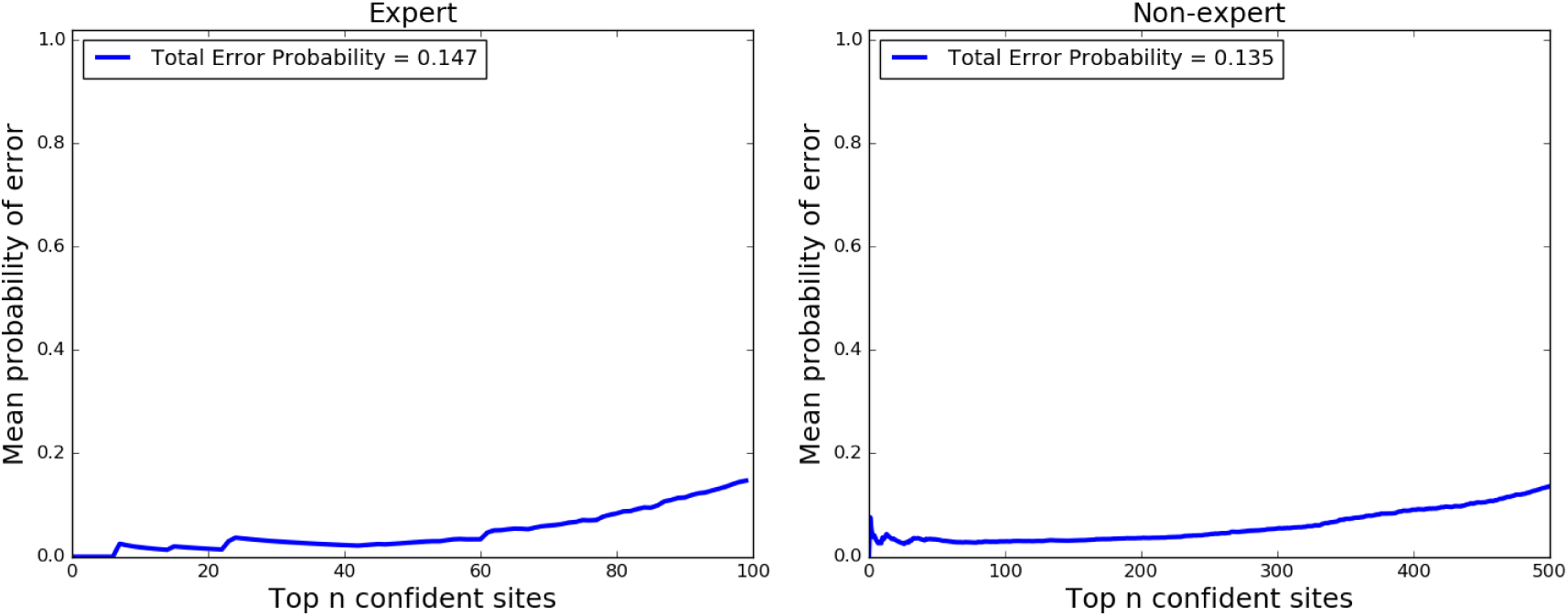
Cumulative mean probability of a Mendelian violation among expert [left] and non-expert [right] classifications as a function of the top n sites. We order sites from highest to lowest score [left to right, x-axis] and for the top n, from 1 to the total number of sites, we compute the average probability of an error for those n sites together. For most confident sites the probability is low and increases as confidence decreases.

**Supplementary Figure 7.**
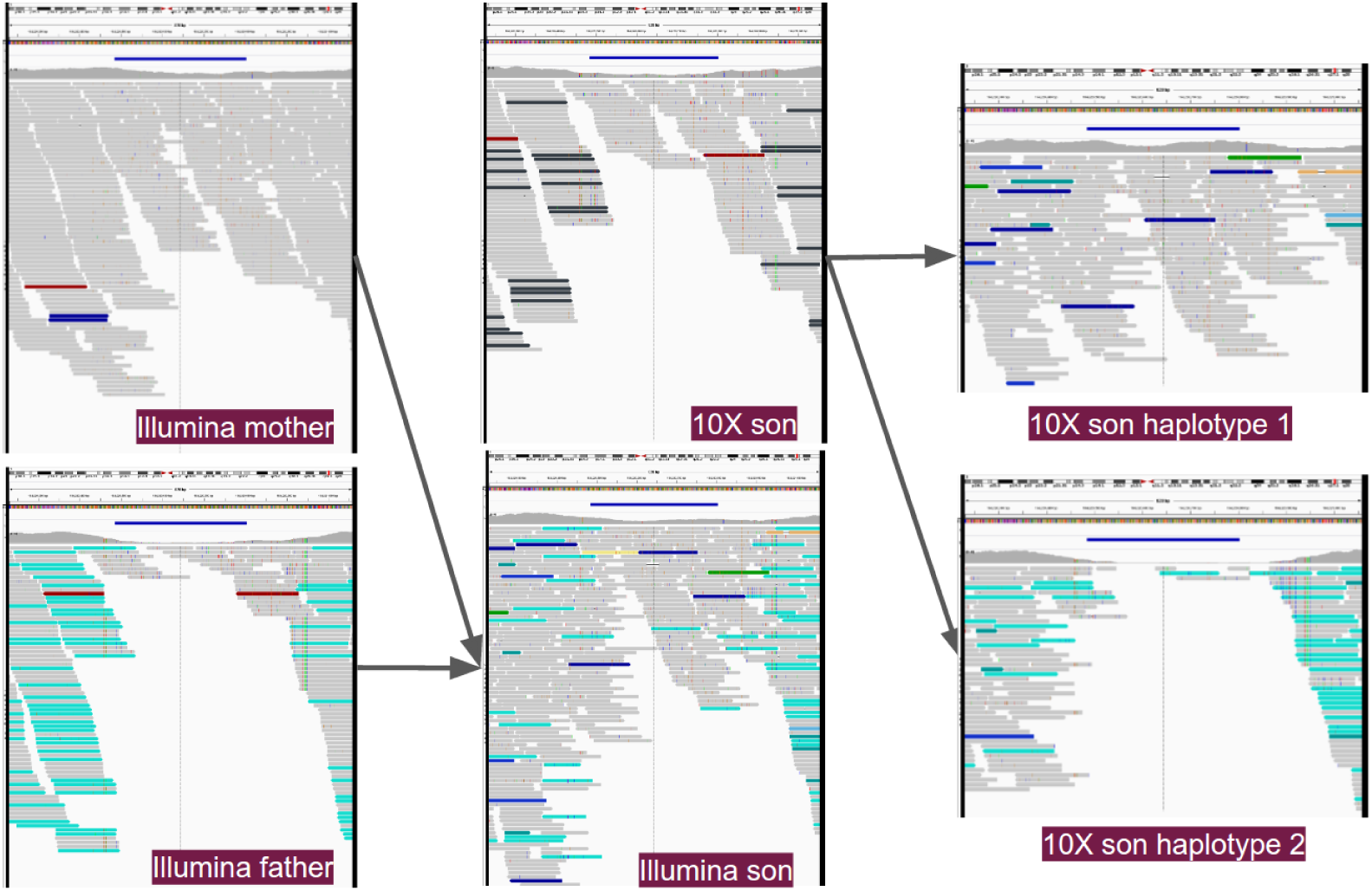
Viewing all image types together shows the power of combining familial and phasing information in different sequencing platforms. This variant (chr3:184220484-184220803) was classified as CN1 in the son with CrowdVariant score 0.93 and is part of the gold set. Svviz classified this example as CN2. Clockwise from top left: Illumina mother, 10X son, 10X son haplotype 1, 10X son haplotype 2, Illumina son, Illumina father. Svviz classification: CN1. Crowd classification: CN2.

**Supplementary Figure 8.**
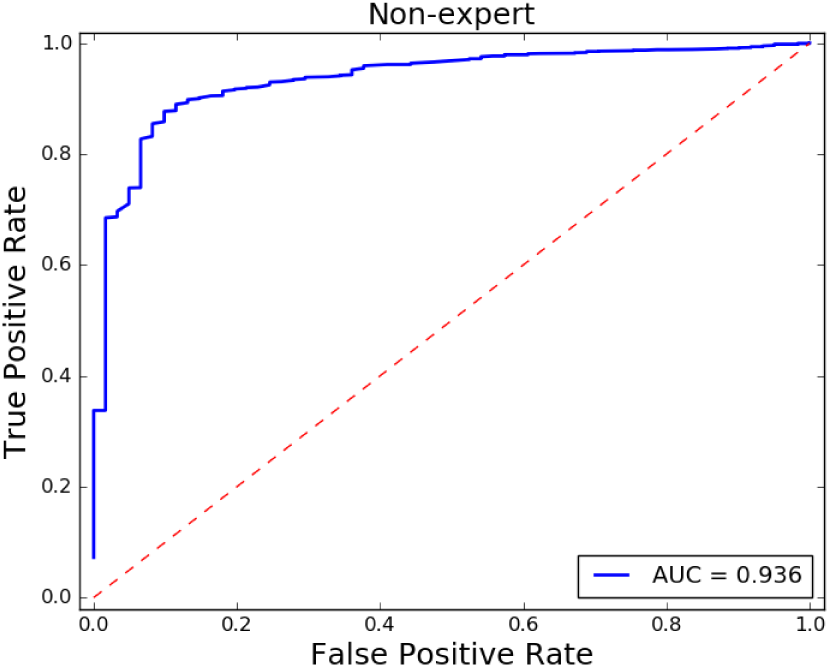
Non-expert genome-wide ROC curves discriminating Mendelian violations (1=no violation, 0=violation) sites by the CrowdVariant score.

**Supplementary Figure 9.**
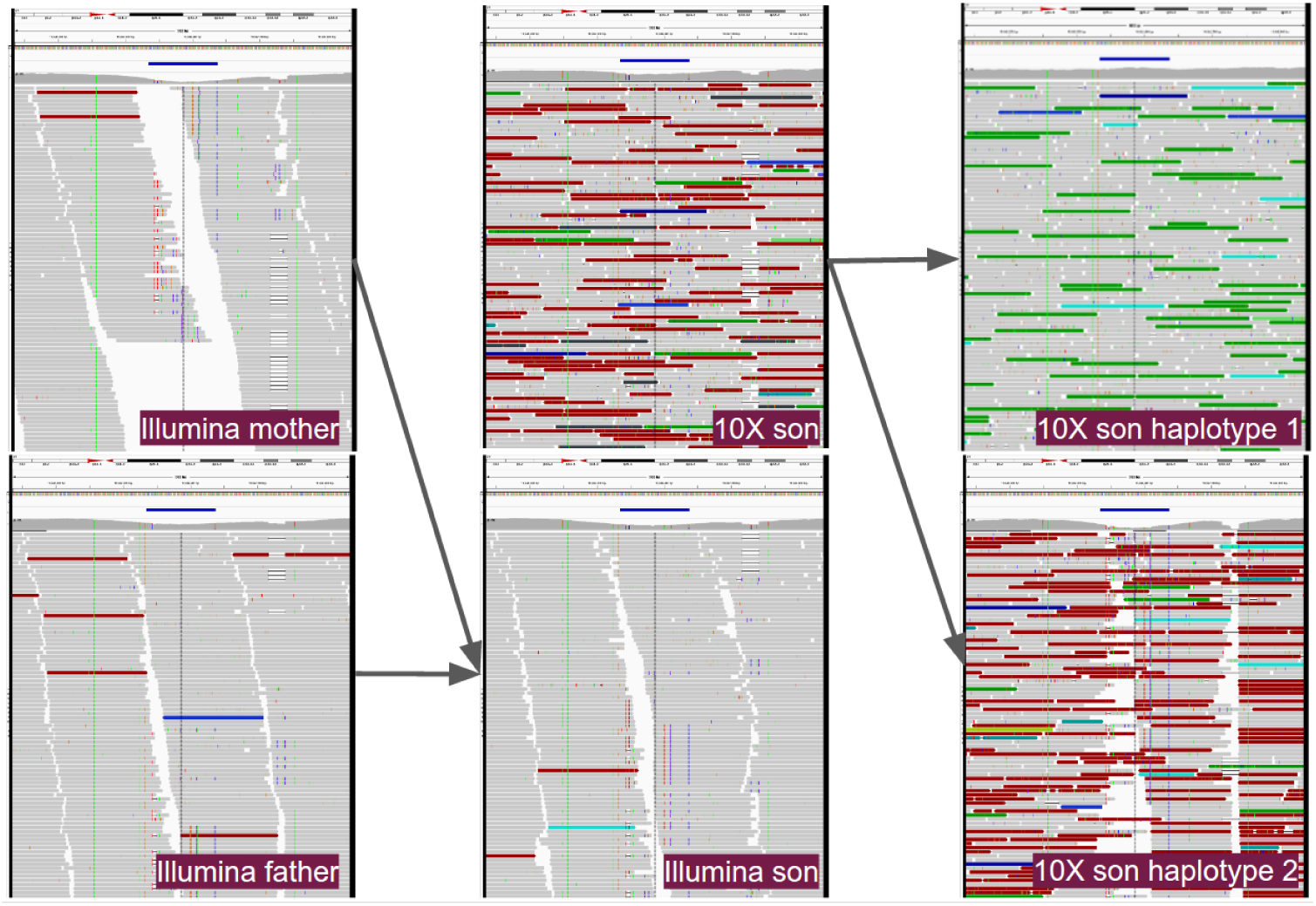
This variant (chr21:10842335-10842437) was the only Mendelian violation in the genome-wide curated set. It was classified as CN2 for both son and father and CN0 for mother. The CrowdVariant score was 0.94.

